# Potentiation of the M_1_ muscarinic acetylcholine receptor normalizes neuronal activation patterns and improves apnea severity in *Mecp2*^+/-^ mice

**DOI:** 10.1101/2024.04.15.586099

**Authors:** Mackenzie Smith, Grace E. Dodis, Amanda M. Vanderplow, Sonia Gonzalez, Yewon Rhee, Rocco G. Gogliotti

**Author notes:** **Declaration of interest:** None.

## Abstract

Rett syndrome (RTT) is a neurodevelopmental disorder that is caused by loss-of-function mutations in the *methyl-CpG binding protein 2* (*MeCP2*) gene. RTT patients experience a myriad of debilitating symptoms, which include respiratory phenotypes that are often associated with lethality. Our previous work established that expression of the M_1_ muscarinic acetylcholine receptor (mAchR) is decreased in RTT autopsy samples, and that potentiation of the M_1_ receptor improves apneas in a mouse model of RTT; however, the population of neurons driving this rescue is unclear. Loss of Mecp2 correlates with excessive neuronal activity in cardiorespiratory nuclei. Since M_1_ is found on cholinergic interneurons, we hypothesized that M_1_-potentiating compounds decrease apnea frequency by tempering brainstem hyperactivity. To test this, *Mecp2*^*+/-*^ and *Mecp2*^*+/+*^ mice were screened for apneas before and after administration of the M_1_ positive allosteric modulator (PAM) VU0453595 (VU595). Brains from the same mice were then imaged for c-Fos, ChAT, and Syto16 using whole-brain light-sheet microscopy to establish genotype and drug-dependent activation patterns that could be correlated with VU595’s efficacy on apneas. The vehicle-treated *Mecp2*^*+/-*^ brain exhibited broad hyperactivity when coupled with the phenotypic prescreen, which was significantly decreased by administration of VU595, particularly in regions known to modulate the activity of respiratory nuclei (i.e. hippocampus and striatum). Further, the extent of apnea rescue in each mouse showed a significant positive correlation with c-Fos expression in non-cholinergic neurons in the striatum, thalamus, dentate gyrus, and within the cholinergic neurons of the brainstem. These results indicate that *Mecp2*^*+/-*^ mice are prone to hyperactivity in brain regions that regulate respiration, which can be normalized through M_1_ potentiation.

## Introduction

RTT is a neurodevelopmental disorder that is caused by *de novo* mutations in a transcription factor known as methyl CpG binding protein 2 (MeCP2) (Amir et al., 1999; Hagberg, 2002). MeCP2 forms a molecular bridge between methylated DNA and chromatin, and mutations that disrupt either interaction are pathogenic (Amir et al., 1999). Clinically, RTT patients experience a period of normal development, followed by stagnation and then rapid regression between 6 and 18 months of age. During this time, acquired social, motor, and cognitive abilities are lost, and symptoms such as breathing irregularities, seizures, and hand-clasping stereotypies develop. These symptoms never resolve and result in severe disability over the patient’s entire life (Feldman et al., 2016; Vashi and Justice, 2019). Current treatment options are limited; however, work from our lab has supported the potential utility of drugs that target the M_1_ muscarinic acetylcholine receptor. Specifically, we demonstrated that M_1_ expression is decreased in the temporal cortex of 40 RTT autopsy samples and that administration of an M_1_ positive allosteric modulator VU0453595 (PAM, VU595) improves social, cognitive, and apnea phenotypes in a *Mecp2*^*+/-*^ mouse model (Smith et al., 2022).

While the M_1_ receptor has a well-defined role in social and cognitive phenotypes (Rusted and Warburton, 1988; Bodick et al., 1997), the mechanisms by which it modifies respiratory patterns are poorly understood. This phenotype is crucial to understand in the context of RTT, where persistent episodes of alternating hyperventilation and breath-holding induce oxidative stress on the autonomic neurons innervating the heart. This pathology is linked to the development of long QT syndrome, which disrupts cardiac rhythm and is responsible for a significant portion of deaths in RTT patients (Bissonnette and Knopp, 2006; Weese-Mayer et al., 2006; Tarquinio et al., 2015; Cordani et al., 2023). The capacity of the M_1_ receptor to rescue apneas in RTT is paradoxical, as the receptor is enriched in the forebrain, but has only minimal expression in the brainstem (Miyoshi et al., 1989; Mallios et al., 1995; Lebois et al., 2018). While the function of excitatory, inhibitory, and modulatory neurons are all compromised within the respiratory nuclei of RTT mice, it is unclear what therapies can be used to normalize activity in these circuits (Ward et al., 2020). One important study found that expression of the immediate early gene c-Fos is increased in brain regions responsible for coordinating the respiratory response to stimuli (i.e. the nucleus tractus solitarus (nTs)) in male *Mecp2*^*-/y*^ models (Kron et al., 2012), indicative of excessive neuronal activity. DREADD based activation of pyramidal neurons in the medial prefrontal cortex (mPFC) reduces apneas in RTT via long-range, inhibitory projections to the locus coeruleus (LC) and its associated connections with respiratory nuclei (Howell et al., 2017). The mPFC regulates behavioral state, and these data align with reports that stress can influence apnea severity (Ren et al., 2012). As M_1_ is both present in brainstem cholinergic interneurons and enriched in the mPFC, there are several potential mechanisms by which M_1_ PAMs correct apneas.

In this study, we combine whole-body plethysmography (WBP) with whole-brain light-sheet imaging to take a non-biased approach to 1) establish genotype-dependent neuronal activation patterns following WBP, 2) identify M_1_ PAM-specific drug signatures throughout the brain, and 3) correlate neuronal activity with the degree of apnea rescue conferred by M_1_potentiation. These experiments established that frontal and cerebellar regions exhibit significant hyperactivity in *Mecp2*^*+/-*^ mice, which is normalized by VU595 administration. Additionally, the number of c-Fos positive neurons in frontal and midbrain regions positively correlates with the degree of apnea rescue conferred by VU595 in *Mecp2*^*+/-*^ mice. Lastly, there is a positive correlation between choline acetyltransferase (ChAT) and c-Fos co-staining in the medulla and the degree of apnea rescue induced by VU595 administration. Together, these data show that the efficacy of M_1_PAMs on respiratory phenotypes is associated with the regulation of non-cholinergic neurons in the fore- and midbrain and cholinergic neurons in the brainstem.

## Materials and Methods

### Animals

Female *Mecp2*^*+/+*^ and *Mecp2*^*+/tm1*.*1 bird*^ mice were purchased from Jackson Laboratories and maintained until experiments were performed at 20 weeks, the age at which reproducible phenotypes are present in our colony (Gogliotti et al., 2017, 2018; Smith et al., 2022). The model and sex used are in line with the standards of the RTT research community, and only female mice were studied to reflect the overwhelmingly female RTT patient population (Katz et al., 2012). In addition, our sample size of n=5/genotype/treatment for each phase of the experiment is in line with what has previously been used to see a significant effect of genotype on c-Fos expression in mouse models of RTT (Kron et al., 2012).

The same n=5 mice from each group were used for both the WBP and tissue collection phases of the experiment to directly correlate the efficacy of VU595 to reduce apneas with the brain regions activated by drug treatment in each mouse. WBP was first performed on mice from all groups to measure apneas before and after VU595 or vehicle administration, followed 90 minutes later by collection of whole-brain samples and downstream light-sheet imaging. All experimental procedures were approved and overseen by the Loyola University Chicago Institutional Animal Care and Use Committee.

### Whole body plethysmography (WBP)

*Mecp2*^*+/-*^ and *Mecp2*^*+/+*^ mice (20w) were placed into a WBP chamber (Buxco, FinePointe) and acclimated for 30 minutes, after which a baseline measure of apneas was recorded for 30 minutes. Mice were then removed from the chamber and administered 10 mg/kg VU595 via intraperitoneal injection or vehicle (10% Tween-80) according to their randomly assigned treatment group, as we have previously established that this dose improves cognitive, social, and respiratory phenotypes in *Mecp2*^*+/-*^ mice at 30-minutes post injection (Smith et al., 2022). Mice were then placed back into WBP chambers, and 30 minutes post-dose apneas were again measured to establish the degree of rescue in each mouse. Apneas, defined as a breath spanning >1 second, were quantified using FinePointe software. *Mecp2*^*+/-*^ mice with an apnea rescue of >10% when administered VU595 were perfused.

### Perfusion

90 minutes post-drug administration, mice were briefly anesthetized with 5% isoflurane and injected with Euthasol. This was followed by transcardial perfusion with heparinized saline and then 4% paraformaldehyde (PFA). Whole brains were harvested and post-fixed in 4% PFA overnight. Samples were stored in PBS+0.02% sodium azide before imaging.

### Light sheet imaging

N=5/genotype/treatment whole mouse brains were processed following the SHIELD protocol (LifeCanvas Technologies, Park et al., 2019). Samples were cleared passively for a week in a delipidation buffer at 45°C. Cleared samples were then actively immunolabeled using SmartBatch+ (LifeCanvas Technologies) based on eFLASH technology integrating stochastic electrotransport (Kim et al., 2015) and SWITCH (Murray et al., 2015). Each brain sample was stained with primary antibody, 24 µL of Syto16 (Thermo Fisher Scientific Inc., S7578), 40 µL of goat anti-ChAT antibody (MilliporeSigma, AB144P), and 3.5 µg of rabbit c-Fos antibody (Abcam, ab214672) followed by fluorescently conjugated secondaries in 1:2 primary: secondary molar ratio (Jackson ImmunoResearch). After active labeling, samples were incubated in EasyIndex (LifeCanvas Technologies) for refractive index matching (RI = 1.52) and imaged at 3.6X magnification, 1.8 µm/pixel resolution on the XY plane, and 4 µm thickness along the z-axis with a SmartSPIM axially swept light sheet microscope (LifeCanvas Technologies).

Sample images were tile-corrected, destriped, and registered to the Allen Brain Atlas (Allen Institute: https://portal.brain-map.org/) using an automated process. A Syto16 channel for each brain was registered to a custom Syto16-based atlas for the Allen Common Coordinate Framework v3, using successive rigid, affine, and b-spline warping algorithms (SimpleElastix: https://simpleelastix.github.io/). Cell locations were found using a custom semantic segmentation model generated using Tensorflow (https://www.tensorflow.org/). Using the atlas registration, measurements were projected onto the Allen Brain Atlas to quantify the ChAT and c-Fos individual cell counts and ChAT/c-Fos co-expression for each atlas-defined region. Representative 3D images of the brain were made in Imaris, and heat maps were made using the average counts in each group in the program Plotly.

### Statistical analysis

Brain regions that had previously been implicated in regulating RTT-like phenotypes were first selected for analysis (Cerebral cortex and hippocampus=Social and cognitive deficits, striatum and midbrain=cognitive and motor deficits, and hindbrain and brainstem=respiration). Next, more detailed brain regions were selected for analysis by filtering for regions with a depth <5 in the Basic Cell Groups and Regions of the Allen Brain Atlas. c-fos and ChAT counts were compared between genotype and treatment groups using 2-way ANOVA and Tukey post-hoc tests. Brain regions were omitted from the analysis of ChAT and c-Fos co-positive counts if ChAT was not expressed in that region. Subjects were removed from light-sheet imaging analysis of a brain region if significant damage occurred to that region during sample collection.

Percent apnea increase was calculated using the formula: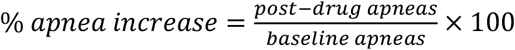, and percent apnea rescue was calculated using the formula: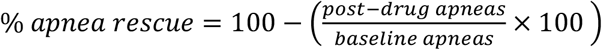.

Correlation analyses were performed using simple linear regression in brain regions implicated in regulating respiration.

## Results

### c-Fos expression is increased in *Mecp2*^*+/-*^ mice and normalized by VU595

Previous work has characterized patterns of hypo- and hyper-activity in the brains of male *Mecp2*^*-/y*^ mice using c-Fos-stained sections (Kron et al., 2012). Here we used a similar approach coupled to whole-brain light sheet imaging to establish baseline and M_1_ PAM activation patterns in female *Mecp2*^*+/-*^ mice. The experimental workflow is outlined in Figure 1A and was performed on 20-week-old *Mecp2*^*+/+*^ and *Mecp2*^*+/-*^ mice. The 20-week timepoint was selected to represent an age where robust phenotypes are reproducibly quantified and responsive to drug treatment within our colony (Gogliotti et al., 2017, 2018; Smith et al., 2022). Control and test mice were 1) habituated to the whole body plethysmograph for 30 min, 2) measured for baseline apnea numbers, 3) administered either the M_1_ PAM VU595 (10mg/kg, ip) or vehicle (10% tween 80), 4) measured for apnea reversal, 5) sacrificed and perfused with PFA, 6) brains were cleared and stained with a pan-neuronal marker (Syto 16), a marker of cholinergic neurons (ChAT), and a maker of neuronal activation (c-Fos), 7) whole-brain light sheet microscopy was performed and patterns of pan-neuronal and cholinergic neuronal activation were mapped to all 840 regions of the Allen Brain Atlas. Representative sagittal, transverse, and coronal images for each of the four test groups are shown in Figure 1B and whole brain 3D videos comparing vehicle-treated *Mecp2*^*+/+*^ and *Mecp2*^*+/-*^ mice (#1), and *Mecp2*^*+/-*^ mice treated with vehicle and VU595 (#2) are provided in Movies 1-2.

**Figure 1:**
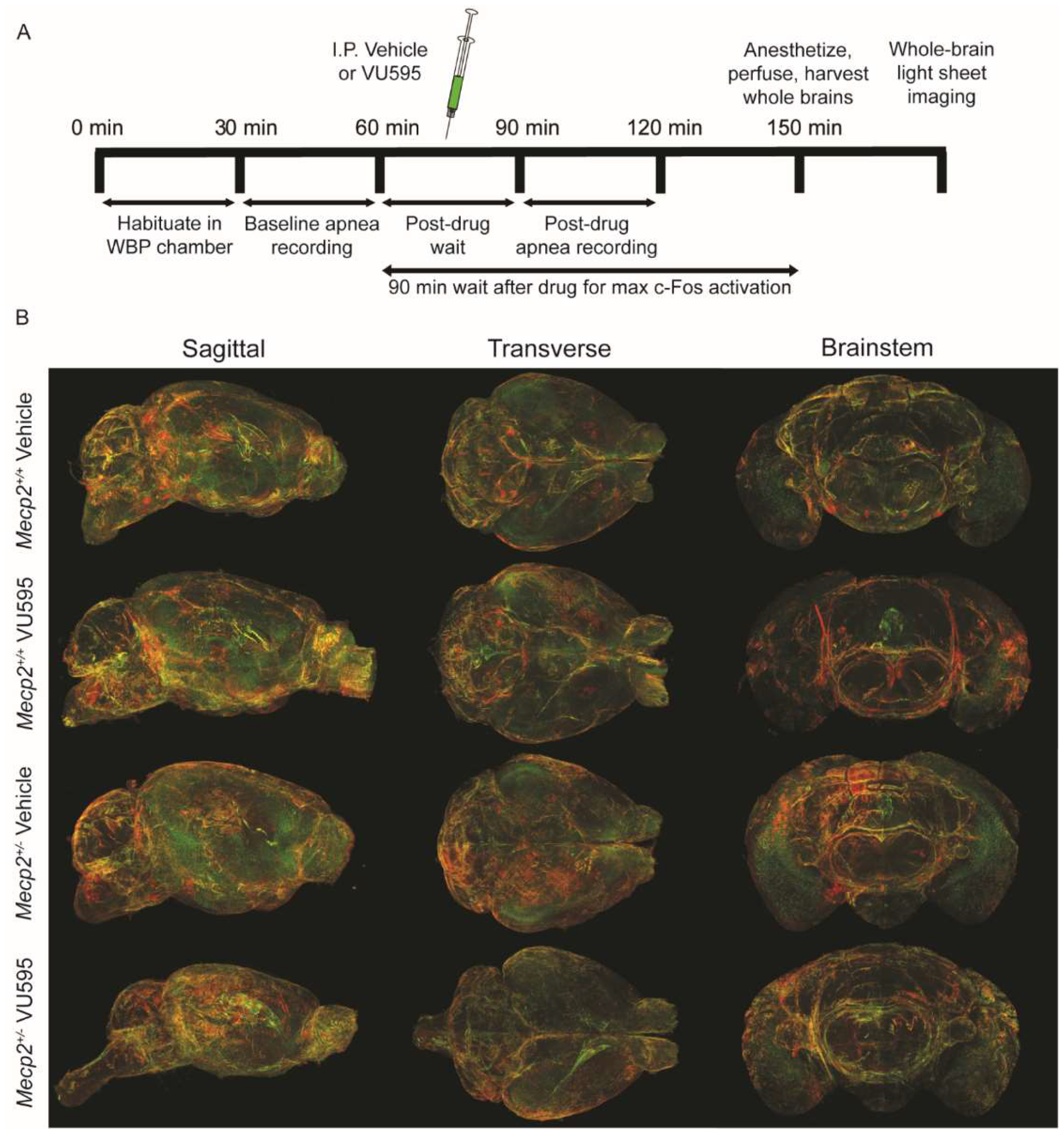
Experimental design and representative images. **A)** Schematic of the protocol used prior to whole brain light sheet imaging. A prescreen was performed to measure apneas in *Mecp2*^*+/+*^ and *Mepc2*^*+/-*^ mice at baseline and after administration of vehicle (10% tween-80) or 10mg/kg VU595. Mice were perfused with PFA 90 minutes after drug administration and whole brains were harvested and post-fixed to prepare for imaging. **B)** Representative whole-brain light sheet sagittal, transverse, and brainstem images. Red = ChAT; Green = c-Fos.

We first used this approach to establish genotype and drug-specific activity signatures in *Mecp2*^*+/-*^ and *Mecp2*^*+/+*^ mice independent of cellular identity. Relative to vehicle-treated *Mecp2*^*+/+*^ mice, heat maps showing average c-Fos counts reveal a consistent pattern of increased activity throughout the vehicle-treated *Mecp2*^*+/-*^ brain, which is reversed by VU595 administration (Fig. 2A). To account for the fact that behavioral state can impact apnea severity, we initially examined a panel of umbrella brain regions implicated in the regulation of several different RTT-like phenotypes (Fig. 2B) (Gabbott et al., 2005; Weese-Mayer et al., 2006; Ren et al., 2012). In this analysis, the number of c-Fos positive neurons in vehicle-treated *Mecp2*^*+/-*^ mice had a broad pattern of hyperactivity (green bars), which reached significance in the hippocampus and striatum and was rescued with VU595 administration (purple bars). No significant differences were found between groups in the cerebral cortex, midbrain, hindbrain, and brainstem.

**Figure 2:**
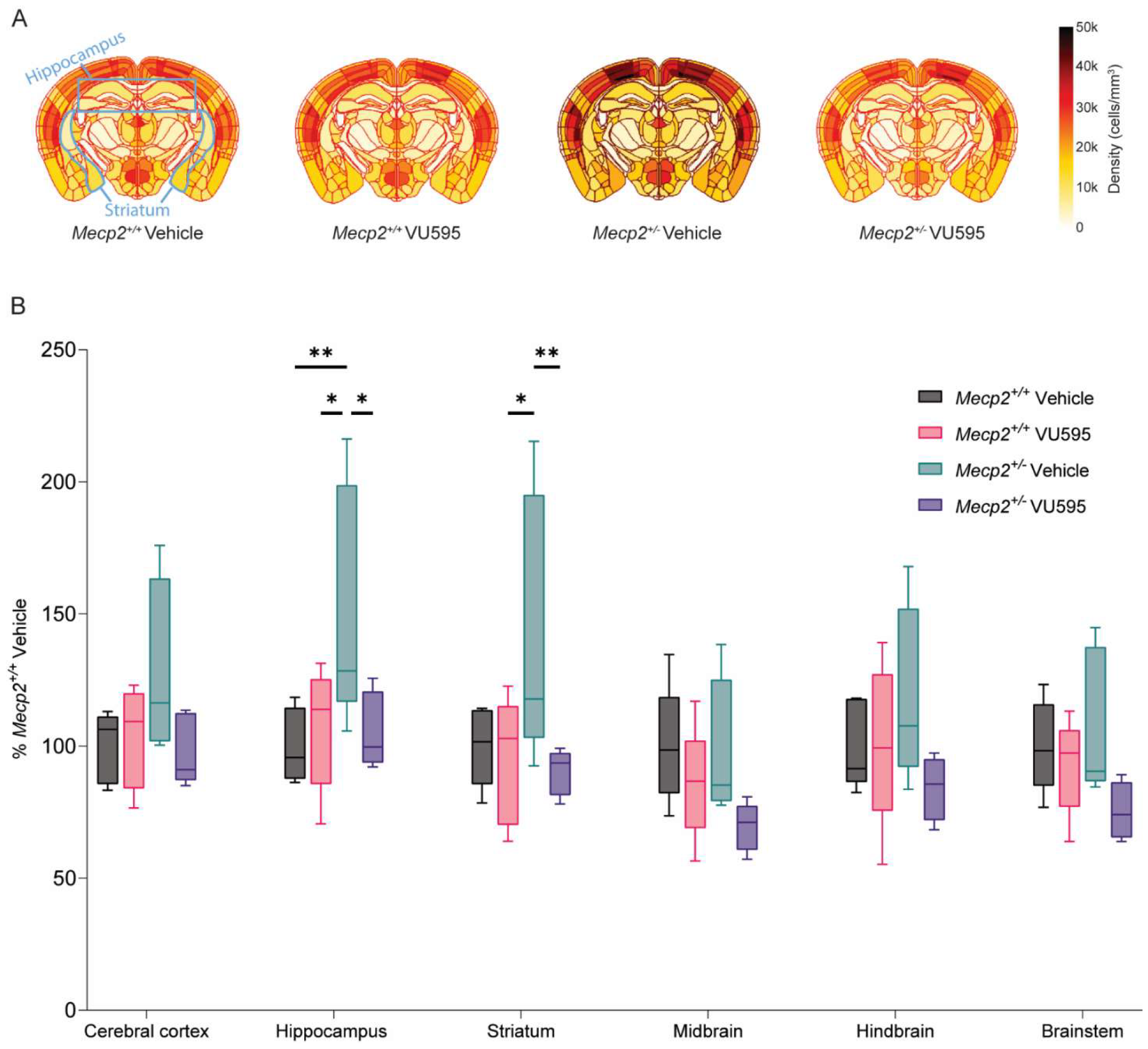
Brain regions associated with RTT-like phenotypes are hyperactive and normalized by M_1_ potentiation in *Mecp2*^*+/-*^ mice. **A)** Heat maps of average c-Fos counts for each group in a representative section of the brain. **B)** Box plots representing c-Fos positive cells normalized to average *Mecp2*^*+/+*^ vehicle in brain regions associated with RTT-like phenotypes. N=4-5/genotype/treatment. Plots span Q1-Q3 with a median centerline. 2-way ANOVA with post-hoc Tukey comparisons test (within each region, compare groups). ^*^p<0.05, ^**^p<0.01. Subject #130 is excluded from hindbrain and brainstem analysis due to tissue damage.

To address the possibility that M_1_ potentiation could be regulating more granular brain regions, we performed an analysis of all divisions nested under “Basic Cell Groups and Regions” in the Allen Brain Atlas with depth <5 parent structures in the tree, which amounted to 28 sub regions. Post-hoc comparisons and significant differences between genotype and/or treatment are listed in Supplemental Table 1, and a comprehensive list of c-Fos activation across the 840 Allen Brain Atlas is provided in Supplemental Table 2. As we anticipated, many of the 28 regions examined showed significant genotype and/or drug effects not represented in the initial umbrella analysis. One example is in the cerebellum, which was not significantly affected when analyzed as an aggregate. With this more focused approach, we quantified a significant increase in c-Fos positive cells in vehicle-treated *Mecp2*^*+/-*^ mice in three cerebellar nuclei (fastigial, interposed, and dentate nucleus), as well as in the vestibulocerebellar nucleus (Fig. 3). VU595 normalized activity patterns to wild type levels in these regions, indicative of a “cooling” effect on neuronal activity that is more pronounced in some sub regions relative to others.

**Figure 3:**
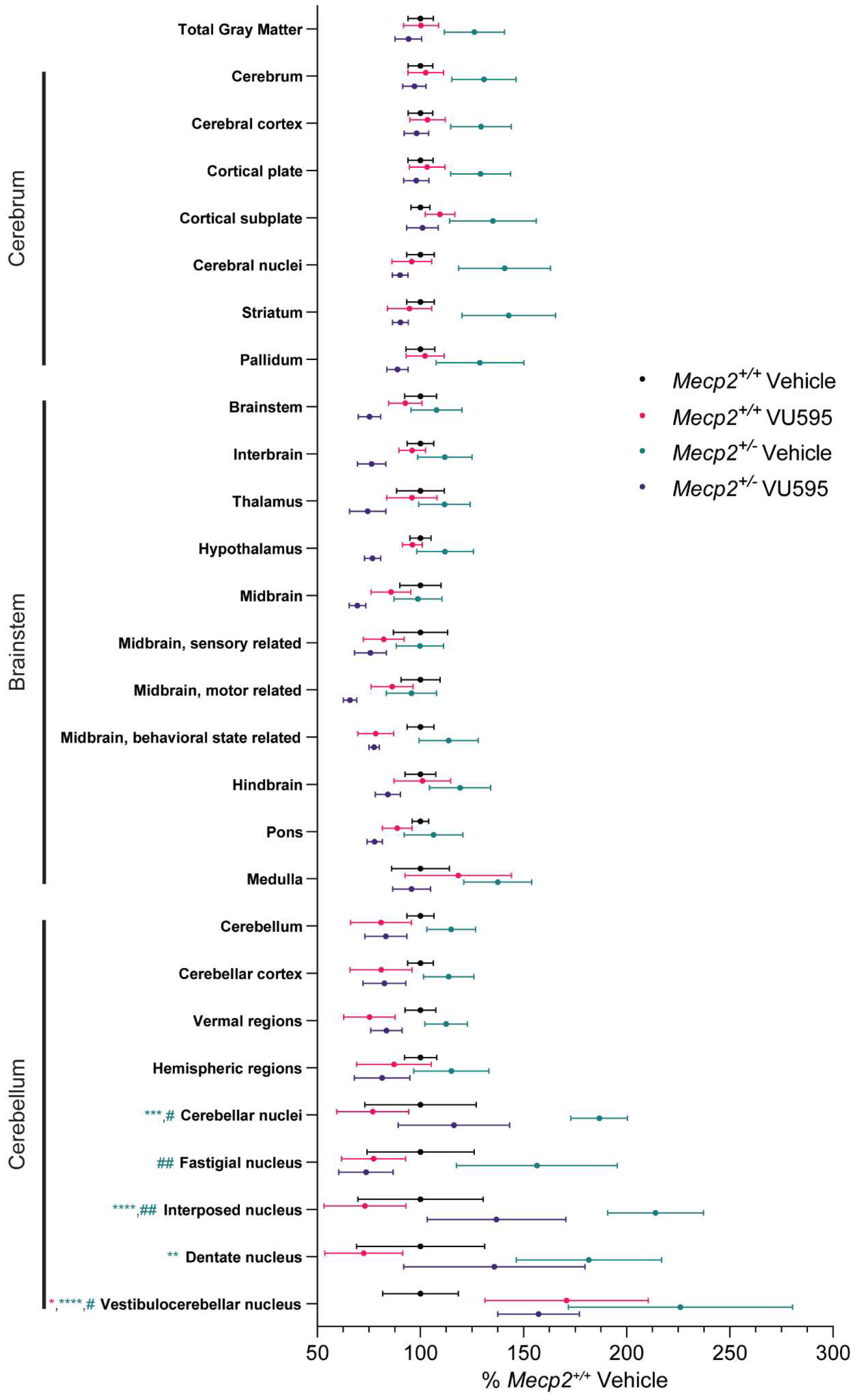
Whole brain effects of VU595 on c-Fos expression. Significant increases in c-Fos expression in *Mecp2*^*+/-*^ mice are seen throughout the brain and normalized by administration of the M_1_ PAM. Basic cell groups and regions with a depth <5 subregions in the Allen Brain Atlas. N=4-5/genotype/treatment. Data shown represent mean ± SEM. 2-way ANOVA with post-hoc Tukey comparisons test. ^***^p<0.001, ^****^p<0.0001 relative to vehicle-treated *Mecp2*^*+/+*^ mice, #p<0.05, # #p<0.01 within genotype comparison. Symbol color denotes the significantly different group. A comprehensive list of statistics and significant comparisons is provided in Supplemental table 1.

### M_1_ PAM-mediated c-Fos activation in cholinergic cells is region-specific

The initial analysis of c-Fos staining in non-specific neurons indicated a general inhibitory effect of M_1_ potentiation, which aligns with the predicted response from activation of cholinergic interneurons. To determine whether this was the case, we next examined the pattern of ChAT and c-Fos co-staining. Heat maps of these results in each experimental group are provided in Fig. 4A. Similar to the previous analysis, the number of c-Fos and ChAT co-positive neurons were first analyzed in umbrella brain regions known to regulate RTT-like behaviors (Fig. 4B). In the hippocampus, the number of c-Fos positive cholinergic neurons was significantly increased in VU595-treated control *Mecp2*^*+/+*^ mice, consistent with known enrichment of the M_1_ receptor in that region (Levey et al., 1995).

**Figure 4:**
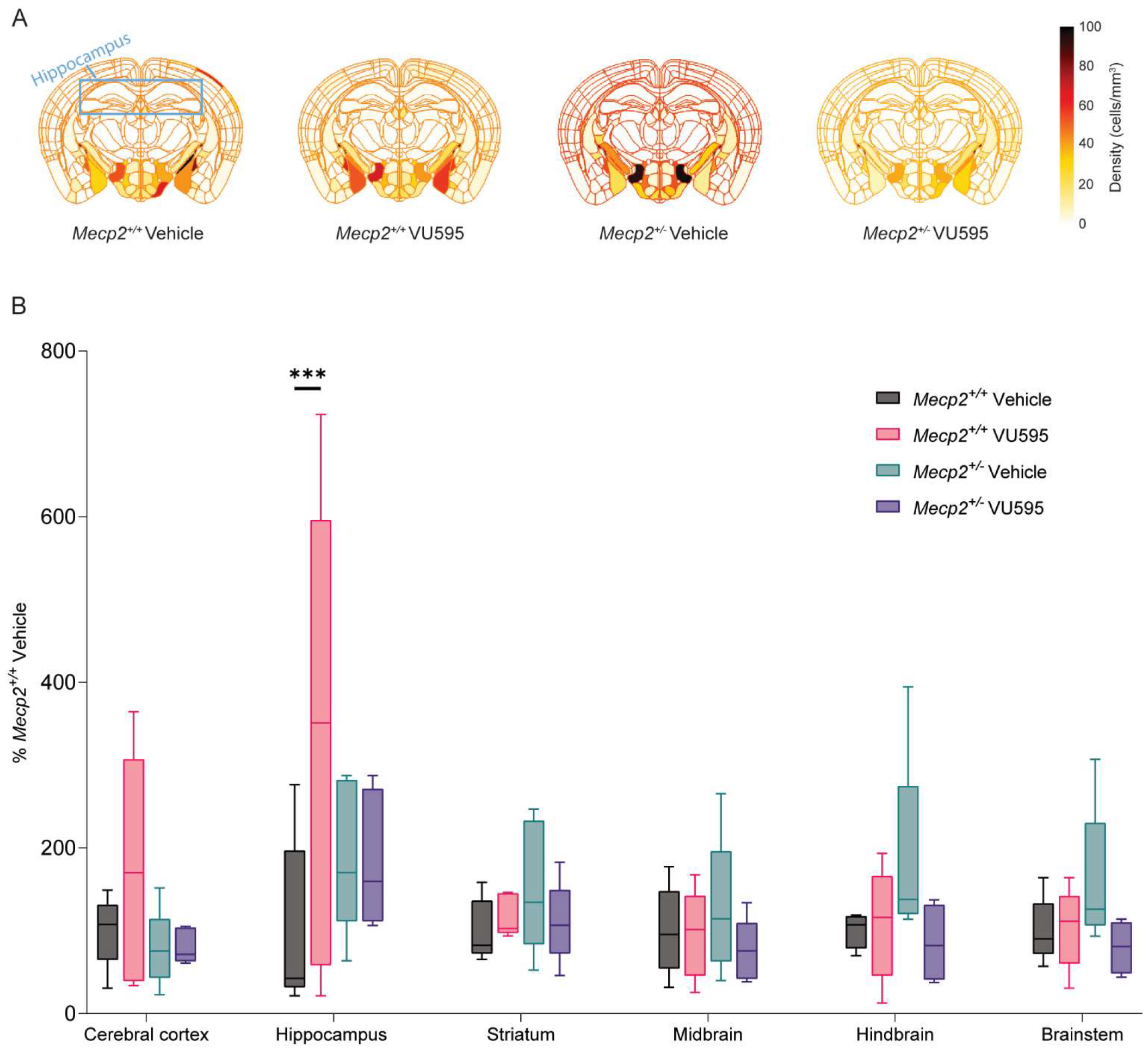
Cholinergic neurons in the hippocampus are activated in *Mecp2*^*+/+*^ administered VU595. **A)** Heat maps of average c-Fos and ChAT co-positive counts for each group in a representative section of the brain. **B)** c-Fos+ChAT counts normalized to average *Mecp2*^*+/+*^ vehicle counts in regions associated with RTT-like phenotypes. N=4-5/genotype/treatment. Plots span Q1-Q3 with a median centerline. 2-way ANOVA with post-hoc Tukey comparisons test. ^***^p<0.001 relative to vehicle-treated Mecp2^+/+^ mice. Subject #130 is excluded from hindbrain and brainstem analysis due to tissue damage.

Three distinct patterns of cholinergic neuronal activation were evident in the comprehensive analysis of 26 “Basic Cell Groups and Regions” examined in the Allen Brain Atlas (fastigial and vestibulocerebellar nuclei were excluded from this analysis due to no ChAT+Fos co-expression in these regions). These patterns show that 1) many regions do not exhibit significant differences between genotypes or treatment groups, including the cerebrum and brainstem; 2) VU595-treated control *Mecp2*^*+/+*^ mice have increased ChAT+c-Fos co-positive staining in cortical, midbrain, and cerebellar regions; and 3) vehicle-treated *Mecp2*^*+/-*^ mice also have increased activity of cholinergic neurons in midbrain and cerebellar regions, which is decreased following administration of VU595 (Fig. 5). When analyzed in concert with the c-Fos alone data, this suggests that both non-specific and cholinergic neurons are hyperactive in the brains of *Mecp2*^*+/-*^ mice. Post-hoc comparisons and significant differences between genotype and/or treatment are listed in Supplemental Table 3, and a comprehensive list of c-Fos and ChAT activation across the Allen Brain Atlas is provided in Supplemental Table 4.

**Figure 5:**
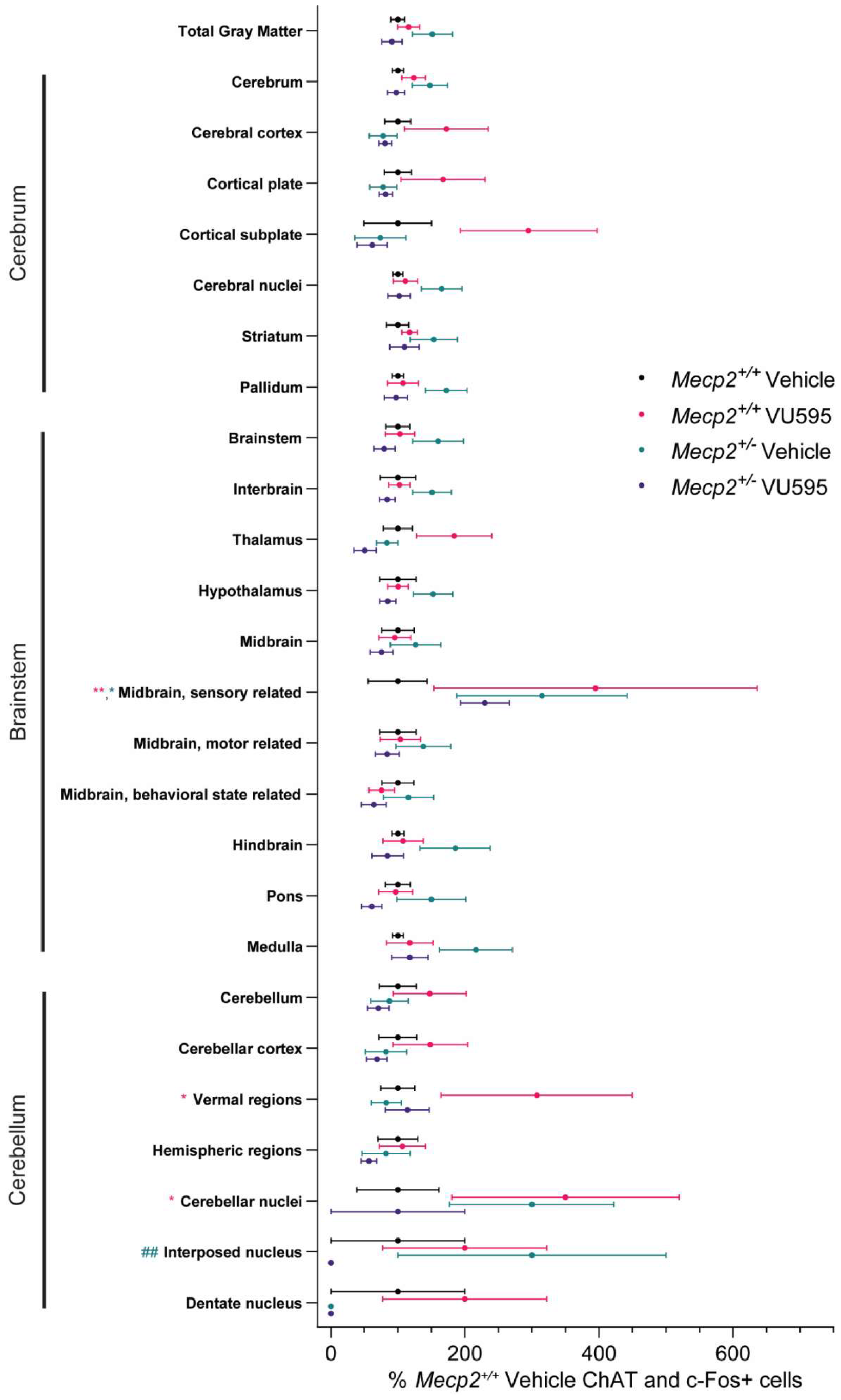
Whole brain effects of VU595 on activation of ChAT positive neurons. *Mecp2*^*+/+*^ and *Mecp2*^*+/-*^ mice present with region specific genotype and drug effects on ChAT positive neuronal activation. Basic cell groups and regions with a depth <5 of the Allen Brain Atlas. N=4-5/genotype/treatment. Data shown represent mean ± SEM. 2-way ANOVA with post-hoc Tukey comparisons test. ^*^p<0.05, ^**^p<0.01, relative to vehicle-treated *Mecp2*^*+/+*^ mice, # #p<0.01 within genotype comparison. Symbol color denotes the significantly different group. A comprehensive list of statistics and significant comparisons is provided in Supplemental table 3.

### Activity in c-Fos+ ChAT neurons correlates with efficacy in reducing apneas

RTT is an X-linked disorder, and female patients and model mice are mosaic for the mutant allele. Consequently, the random distribution pattern of the mutant allele can lead to heterogeneity in disease presentation and treatment response. To account for variability, we performed a phenotypic pre-screen of the mice used to generate Figures 2-5 to establish the percentage of M_1_-PAM-dependent apnea reversal in each individual mouse, which could then be correlated with the magnitude of c-Fos activation across their brain. The number of apneas at baseline and following drug treatment is shown in Figure 6A-D. In line with our previous findings (Smith et al., 2022), *Mecp2*^*+/-*^ mice who were administered VU595 showed a significant reduction in apnea number. Conversely, vehicle-treated *Mecp2*^*+/-*^ mice had significantly more apneas relative to baseline, potentially indicative of increased anxiety associated with the test.

**Figure 6:**
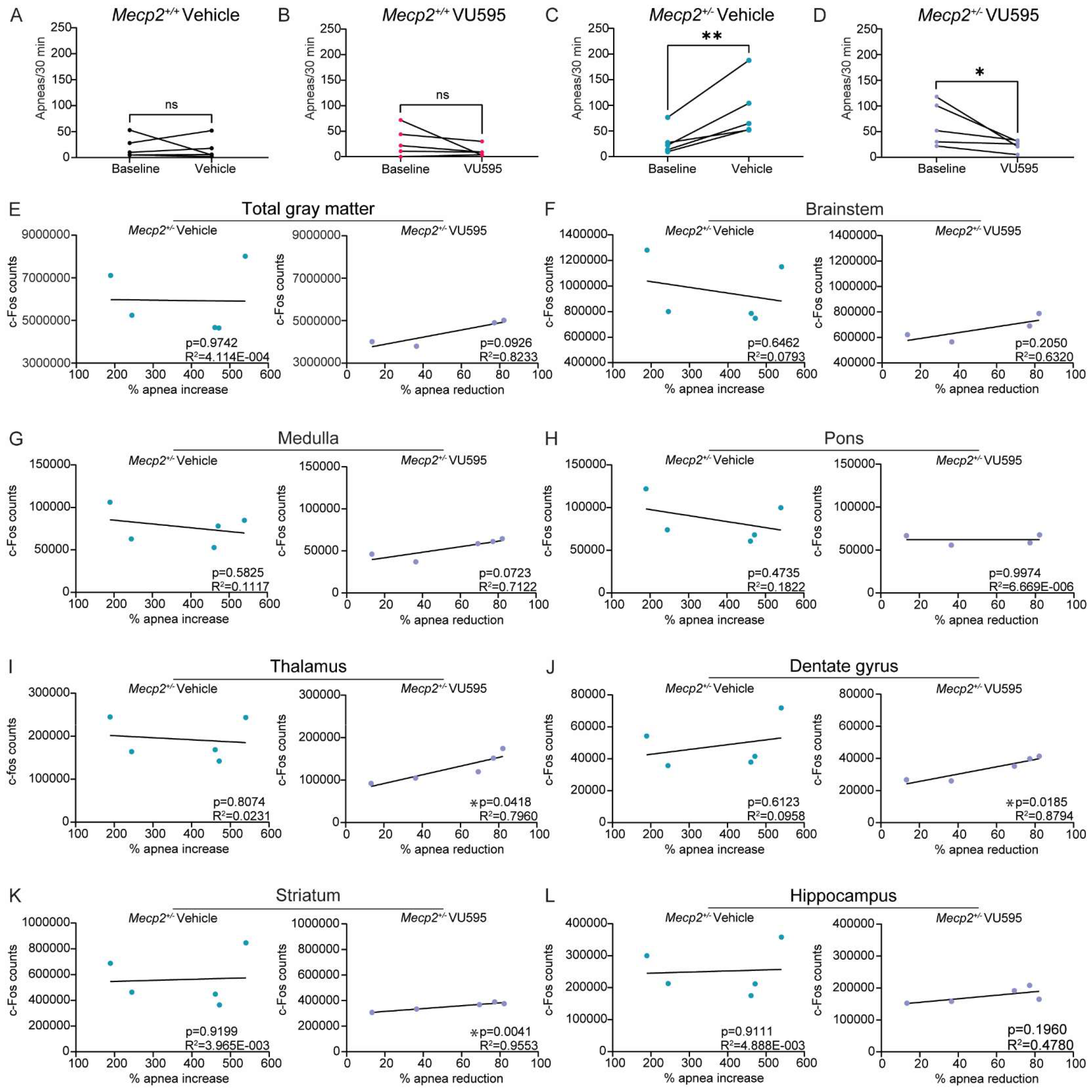
The activity of fore- and midbrain neurons is involved in regulating apneas. **A-D)** Whole body plethysmography. N=5/genotype/treatment. T-test. **A-B)** The respiratory pattern of control *Mecp2*^*+/+*^ mice is unaffected by vehicle and/or VU595 treatment. **C)** Apneas are significantly increased in vehicle-treated *Mecp2*^*+/-*^ mice relative to baseline recording. **D)** Consistent with our previous reports, apneas were significantly reduced in *Mecp2*^*+/-*^ mice following administration of VU595. **E-L)** Correlation between c-Fos positive cell counts and % apnea reduction relative to baseline in vehicle- (left) or VU595- (right) treated *Mecp2*^*+/-*^ mice. Significant correlations were quantified in the thalamus, dentate gyrus, and striatum in the VU595-treated group. N=4-5/genotype/treatment. Simple linear regression. Subject 130 is excluded from the total gray matter, brainstem, and pons due to significant damage in these regions.

We initially compared c-Fos expression relative to percent apnea increase (vehicle) or decrease (VU595) in *Mecp2*^*+/-*^ mice independent of cellular identity. This analysis showed no significant correlations in total gray matter or brain regions containing respiratory nuclei in either group (Fig. 6E-H, brainstem region, medulla, pons). In contrast, we quantified a significant correlation between the number of c-Fos positive neurons activated by VU595 administration and the degree of apnea reversal in brain regions that project to respiratory nuclei, either directly or indirectly. These regions include the thalamus, dentate gyrus, and striatum (Fig. 6 I-L). No significant correlations between apnea increase and c-Fos expression were quantified in vehicle-treated *Mecp2*^*+/-*^ mice.

We next compared the magnitude of c-Fos activation in ChAT-positive neurons relative to percent apnea increase (vehicle) or decrease (VU595) in *Mecp2*^*+/-*^ mice. Overall, there was no correlation between apnea increase and cholinergic activity in the vehicle control *Mecp2*^*+/-*^ group (Fig. 7). In direct contrast to what was observed with c-Fos alone, VU595-treated *Mecp2*^*+/-*^ mice showed a significant correlation between the percentage of apnea rescue and the number of activated cholinergic neurons in the brainstem and medulla (Fig. 7 A-C), where respiratory nuclei are located. No significant correlation was seen in the pons (Fig. 7D). Again, in opposition to c-Fos alone, we did not quantify significant correlations in regions that project to respiratory nuclei (Fig. 7E-G, thalamus, dentate gyrus, and striatum). Interestingly, we observed a significant negative correlation in the hippocampus, where less cholinergic activation was associated with a greater decrease in apneas (Fig. 7H). Together, these data may suggest that M_1_ activation in regions outside of respiratory nuclei stimulates cholinergic neurons within the brainstem to enhance inhibitory tone and normalize hyperactivity in *Mecp2*^*+/-*^ mice.

**Figure 7:**
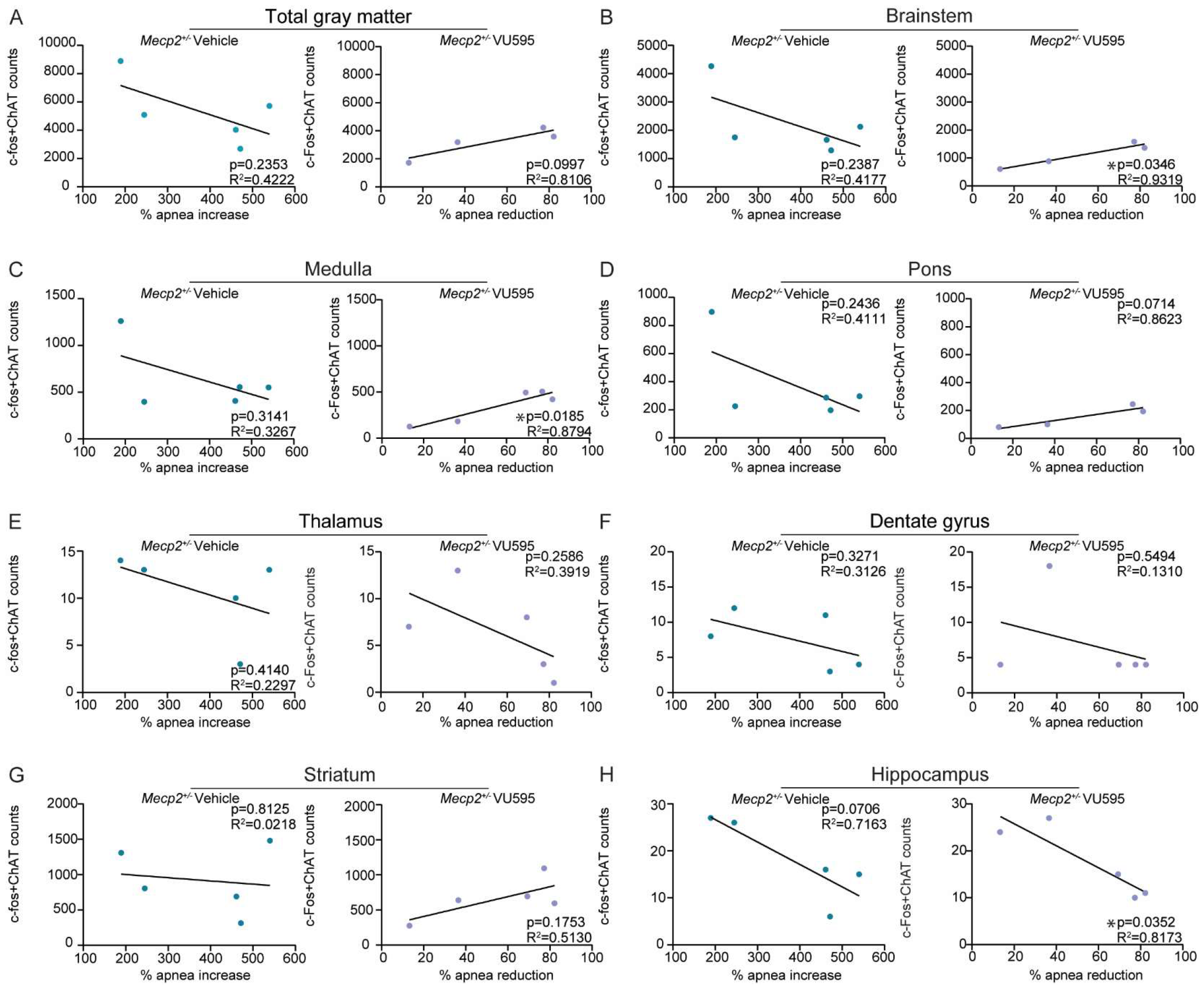
The activity of brainstem cholinergic neurons correlates with apnea rescue. N=4-5/genotype/treatment. **A-H)** Correlation between c-Fos and ChAT co-positive cell counts and % apnea reduction relative to baseline in vehicle- (left) or VU595 (right)-treated *Mecp2*^*+/-*^ mice. Significant correlations are seen in the brainstem, medulla, and hippocampus (hippocampal region) in the VU595-treated group. Linear regression analysis. Subject 130 is excluded from the total gray matter, brainstem, and pons due to significant damage in these regions. N=4-5/genotype/treatment.

## Discussion

Respiratory phenotypes in RTT patients and model mice are state-dependent, where factors like anxiety can impact the severity and prevalence of apneas (Weese-Mayer et al., 2006; Ren et al., 2012). Consistent with this theory, our data demonstrate a pattern of increased expression of the immediate early gene c-Fos throughout the brain of *Mecp2*^*+/-*^ mice that have undergone a phenotypic pre-screen, which contrasts with published reports from mice harvested at baseline (Kron et al., 2012). Our results also show that the activation of non-cholinergic neurons in frontal and midbrain regions and cholinergic neurons of the brainstem correlate with the magnitude of M_1_ PAM-dependent apnea rescue in RTT model mice. These studies provide a blueprint for how drug efficacy can be correlated with neuronal activation patterns in a non-biased manner using whole-brain light sheet imaging.

Our initial analyses quantified a significant increase in c-Fos positive neurons in the hippocampus and striatum of *Mecp2*^*+/-*^ mice, which was both rescued by VU595 administration and correlated with a decrease in apneas. This finding supports reports that the hippocampus is prone to hyper-excitability following a challenge with excitatory stimuli in RTT (Zhang et al., 2008). While classically implicated in cognition, the role of the hippocampus in mediating respiration via the prefronto-thalamic network has been documented (Basha et al., 2023). In this circuit, respiratory rhythm is sensed by thalamic nuclei, which then act as a bridge between the PFC and hippocampus to promote neuronal synchrony required for the cognitive operations performed by these regions. In addition, the striatum and hippocampus also function in response to stress (Jacobson and Sapolsky, 1991; Plattner et al., 2015). The influence of the behavioral state on cardiorespiratory symptoms in patients with RTT bolsters the hypothesis for a behavioral component in mediating the presence and severity of apneas, which could potentially be corrected by normalizing activity in these regions (Weese-Mayer et al., 2008). Additionally, these results support our previous manuscript, which demonstrated that M_1_ potentiation rescues not only apneas but also social and cognitive behaviors that are linked to hippocampal regulation (Smith et al., 2022).

One unexpected result was that VU595 administration significantly decreased c-Fos expression in the cerebellum of *Mecp2*^*+/-*^ mice, as this brain region is not commonly implicated in phenotypes rescued by M_1_ PAMs. However, altered gene expression, including decreased M_1_ receptor levels, has been observed in human post-mortem cerebellar samples from RTT patients (Gogliotti et al., 2018) and our data further suggest a role for M_1_ signaling in that brain region. Additionally, projections from the cerebellum target respiratory nuclei such as the Kölliker-Fuse nucleus and locus coeruleus, and activity in the cerebellum is increased during respiratory challenge (Teune et al., 2000; Critchley et al., 2015; Schwarz et al., 2015; Fujita et al., 2020). Together, this provides several routes by which disrupted cerebellar activity in RTT is linked to M_1_ activity and respiration.

The effects of VU595 on cholinergic neurons were assessed using c-Fos+ChAT co-staining. As expected, there was increased activity in cholinergic neurons in *Mecp2*^*+/+*^ control mice administered VU595 in areas of the brain where M_1_ expression is enriched (i.e. hippocampus). Similar to what was found with cell type agnostic c-Fos staining, we also noted significant effects of genotype and treatment on c-Fos and ChAT co-positive staining in the cerebellum, further highlighting the capacity of M_1_to modulate neuronal activity in that region.

Among our most important findings was the positive correlation between the degree of apnea rescue with the activity of 1) non-specific neurons in frontal and midbrain regions and 2) cholinergic neurons of the brainstem. This could suggest that excitatory neurons in frontal regions project to cholinergic neurons in the brainstem, thereby increasing inhibitory tone on hyperactive respiratory nuclei. Such a context could explain how M_1_ potentiation resolves apneas, given that it is primarily enriched in frontal regions (Lebois et al., 2018). It is paradoxical that increased cholinergic activation correlates with apnea rescue, given that the pattern of hyperactivity is independent of cell type *Mecp2*^*+/-*^ mice. While speculative, this may indicate that the reduction of excitatory tone and the preservation of cholinergic tone is crucial to improving apneas in RTT. These results could also point to a smaller subset of cholinergic interneurons within the medulla, which regulate respiratory circuits directly (Miyoshi et al., 1989; Mallios et al., 1995). Further work is required to tease apart the mechanistic basis of this correlation.

Our experiments largely agree with what is currently known about the excitatory-inhibitory imbalance in RTT, particularly with regard to inhibitory neurotransmission. Removing MeCP2 from inhibitory neurons produces RTT-like phenotypes in mice, which are rescued when MeCP2 levels are restored in this cell type (Chao et al., 2010; Ure et al., 2016). Moreover, enhancing GABAergic signaling improves respiratory phenotypes (Abdala et al., 2010; Johnson et al., 2020; Wu et al., 2021). These findings suggest that a global lack of inhibitory tone plays a role in respiratory dysfunction in RTT. Our results showing that cholinergic activation within the medulla correlates with reduced apneas further support this view.

We acknowledge that the hyperactivity phenotype reported in fore and midbrain structures here is in direct contrast to the hypoactivity phenotype described in these regions by others (Kron et al., 2012). However, several key differences in experimental design could account for this discrepancy. The most critical is likely that historical studies have been performed at baseline, whereas our experiments involved handling, injection, and WBP steps that may induce stress. It has been shown that respiratory abnormalities are correlated with blood corticosterone levels in RTT (Ren et al., 2012), and therefore our findings may more directly correlate with state-dependent respiratory phenotypes as opposed to apneas at rest. This may also explain why vehicle-treated *Mecp2*^*+/-*^ mice have enhanced apneas relative to their baseline in our studies (Fig. 6C). MeCP2 is an activity-dependent gene regulator, and it is known that excitatory neurons in regions like the hippocampus maintain a prolonged state of activation following stimulation (Li et al., 2016). Consequently, the increase in c-Fos may be representative of an inability of activated neurons to reset patterns of immediate early gene expression. A second important variable is that the *Mecp2*^*+/-*^ mice used in our experiments were older than those in previous reports. This potentially indicates that the landscape of inhibitory and excitatory disruptions may change with age and disease progression. Continued work in this area will be critical to determining the extent to which these and other variables contribute to the results presented herein.

Currently, there is only one FDA-approved drug for RTT, and our results have implications for future avenues for the development of therapeutics for this disorder. Pharmacological activation of the cholinergic system has previously shown efficacy in clinical trials for schizophrenia and Alzheimer’s disease in treating social and cognitive deficits, which are two symptom domains that are also disrupted in RTT (Bodick et al., 1997; Sauder et al., 2022). This research suggests the potential for repurposing these medications for RTT. Beyond RTT, we introduce a methodology for establishing drug and genotype signatures in whole-brain samples, with broad applicability in neuropharmacology. This approach can be used to provide a non-biased view of region-specific effects of therapeutics to help determine the population of cells driving the rescue of a phenotype, or even identify the involvement of unexpected circuits or regions. By employing this method to characterize the action of the M_1_ PAM VU595 in the brain, we show that potentiation of the M_1_ receptor corrects hyperactivity in RTT. Further, the effect of M_1_ potentiation on apneas is driven by the regulation of activity in both non-cholinergic projections and cholinergic neurons of the brainstem. These results advocate for the use of M_1_ PAMs as well as other compounds that temper excitability in the brain as possible treatments for RTT.

## Acknowledgments

The authors acknowledge and thank Dr. Gwendolyn Kartje and Brian Powers for technical instruction and resource sharing. The authors would also like to thank Dr. Colleen Niswender for her intellectual contributions to project planning and manuscript preparation.

